## Supplementary File 2 for "Genetic basis of *Arabidopsis thaliana* responses to infection by naïve and adapted isolates of turnip mosaic virus"

| **Table S1.** Genetic associations in the GWA analysis of the ancestral and evolved TuMV isolates including all loci. Table shows the most strongly associated SNP in each peak, and annotated genes within the region spanned by significantly associated SNPs. | | | | | | |
| --- | --- | --- | --- | --- | --- | --- |
| **Chromosome** | **Position** | **−log_10_*P*** | **Allele frequency** | **Effect** | **Genes** | **Function** |
| 1 | 6,167,132 | 10.603 | 0.036 | isolate-specific | *MAIN* | Meristem development and transposon silencing |
|  |  |  |  |  | *AT1G17940* | Endosomal targeting |
| 1 | 12,086,029 | 8.511 | 0.031 | isolate-specific | *AT1G33320* | Methionine biosynthesis |
|  |  |  |  |  | *AT1G33330* | RNA metabolism and translation release |
|  |  |  |  |  | *FAS4* | Helicase activity, regulation of RNA metabolism |
| 1 | 21,740,662 | 8.119 | 0.031 | common | *RXW8* | GDSL-like lipase/acylhydrolase; cell differentiation |
|  |  |  |  |  | *RPP7* | LRR disease-resistance protein |
| 1 | 22,508,950 | 9.534 | 0.031 | isolate-specific | *AT1G61100* | TIR disease-resistance protein |
| 2 | 2,348,583 | 8.146 | 0.033 | common | *GNAT6* | Acyl-CoA N-acetyltransferase; protein modification |
|  |  |  |  |  | *RAD7A* | Nucleotide-excision repair, response to UV-C radiation |
| 2 | 2,945,626 | 8.551 | 0.033 | common | *CAS1* | Cycloartenol synthase; pollen development |
|  |  |  |  |  | *AT2G07120* | F-box associated ubiquitination |
| 2 | 5,927,469 | 74.278 | 0.034 | common | *ALD1* | AGD2-like defense protein, systemic acquired resistance |
|  |  |  |  |  | *AT2G14070* | Wound repair |
|  |  |  |  |  | *AT2G14080* | TIR-LRR-NBS disease-resistance protein |
|  |  |  |  |  | *CALS5* | Callose synthesis |
|  |  |  |  |  | *CYP705A13* | Cytochrome P450 |
|  |  |  |  |  | *AT2G14110* | Haloacid dehalogenase-like hydrolase; protein modification |
|  |  |  |  |  | *DRP3B* | Membranes fission and fusion, viral replication |
| 2 | 6,763,787 | 11.198 | 0.033 | common | *AT2G15500* | RNA binding |
|  |  |  |  |  | *CTL03* | Protein ubiquitination; meristem development |
|  |  |  |  |  | *FDA9* | F-Box family protein |
| 2 | 8,198,266 | 8.115 | 0.045 | isolate-specific | *AT2G18920* | Hypothetical protein |
|  |  |  |  |  | *AT2G18938* | Transmembrane protein |
| 3 | 10,663,084 | 9.721 | 0.031 | isolate-specific | *GFS9* | Membrane trafficking in the Golgi apparatus |
|  |  |  |  |  | *BIR2* | LRR negative regulator of pattern-triggered immunity |
| 3 | 12,668,171 | 10.388 | 0.039 | common | *AT3G31005* | Pseudogene of *AT1G66520* (*PDE194*) |
| 4 | 3,920,265 | 10.467 | 0.039 | isolate-specific | *AT4G06688* | Myosin-heavy-chain-like protein |
| 5 | 10,050,611 | 7.782 | 0.093 | common | *AT5G28053* | LINE transposon element |
| 5 | 16,238,180 | 8.597 | 0.035 | isolate-specific | *AT5G40540* | Protein kinase superfamily protein; signal transduction |

| **Table S2.** Genetic associations in the GWA analysis of the ancestral and evolved TuMV isolates that included the SNP at Chr2:5927469 as a covariate. Table shows the most strongly associated SNP in each peak, and annotated genes within the region spanned by significantly associated SNPs. | | | | | | |
| --- | --- | --- | --- | --- | --- | --- |
| **Chromosome** | **Position** | **−log_10_*P*** | **Allele frequency** | **Effect** | **Genes** | **Function** |
| 1 | 6,167,132 | 11.030 | 0.036 | isolate-specific | *MAIN* | Meristem development and transposon silencing |
|  |  |  |  |  | *AT1G17940* | Endosomal targeting |
| 1 | 12,086,029 | 8.686 | 0.031 | isolate-specific | *AT1G33320* | Methionine biosynthesis |
|  |  |  |  |  | *AT1G33330* | RNA metabolism and translation release |
|  |  |  |  |  | *FAS4* | Helicase activity, regulation of RNA metabolism |
| 1 | 22,508,950 | 9.567 | 0.031 | isolate-specific | *AT1G61100* | TIR disease-resistance protein |
| 2 | 8,198,266 | 8.624 | 0.045 | isolate-specific | *AT2G18920* | Hypothetical protein |
|  |  |  |  |  | *AT2G18938* | Transmembrane protein |
| 3 | 10,663,084 | 10.323 | 0.031 | isolate-specific | *GFS9* | Golgi apparatus |
|  |  |  |  |  | *BIR2* | LRR negative regulator of pattern-triggered immunity |
| 4 | 273,465 | 8.121 | 0.170 | isolate-specific | *FRIGIDA* | Negative regulator of flowering development |
|  |  |  |  |  | *AT4G00651* | Iron superoxide dismutase; protein modification |
|  |  |  |  |  | *RH8* | RNA helicase; intracellular transport of viral proteins |
| 4 | 3,920,265 | 9.994 | 0.039 | isolate-specific | *AT4G00668* | Myosin-heavy-chain-like protein |
| 4 | 16,556,249 | 9.155 | 0.032 | common | *AT4G34690* | Transmembrane protein |
|  |  |  |  |  | *CIB22* | Mitochondrial subunit; photorespiration |
|  |  |  |  |  | *ADC2* | Arginine decarboxylase; response to ABA and JA |
| 5 | 2,386,447 | 7.976 | 0.037 | common | *GRP16* | Flower development |
| 5 | 16,238,180 | 8.236 | 0.035 | isolate-specific | *AT5G40540* | Protein kinase superfamily protein; signal transduction |
