## Supplementary material for "Genetic basis of *Arabidopsis thaliana* responses to infection by naïve and adapted isolates of turnip mosaic virus": Figure 2 - figure supplement 1

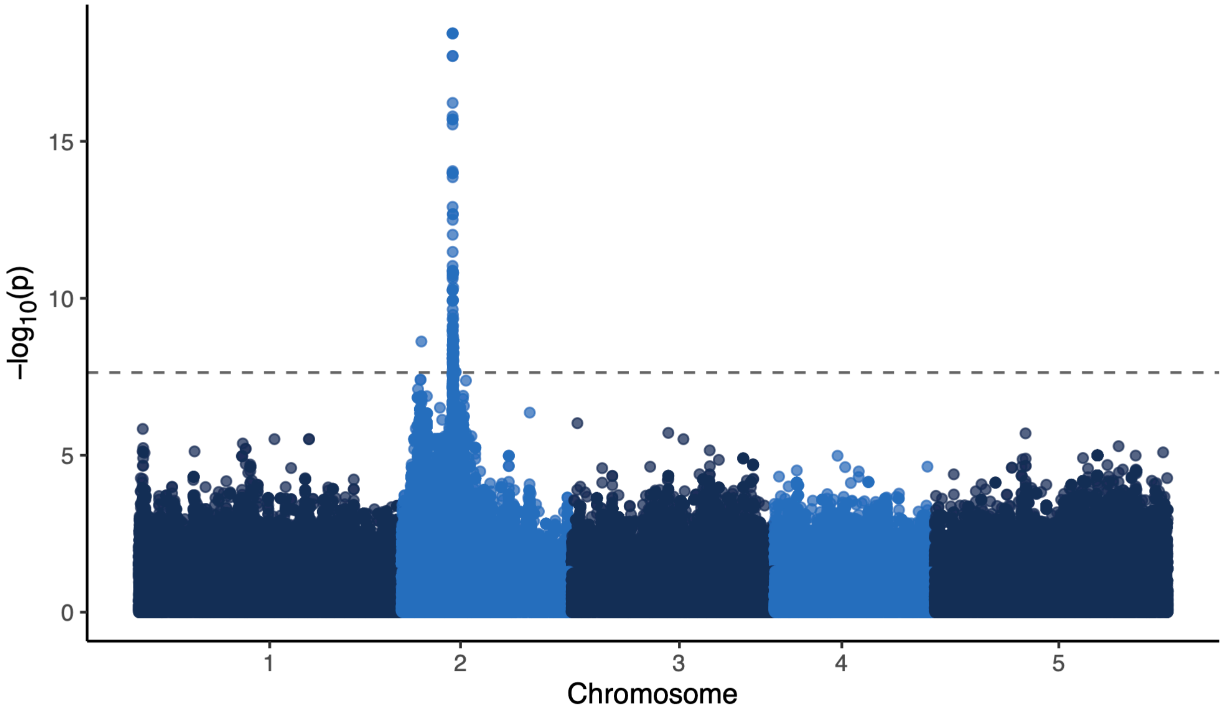


Figure S1: Manhattan plots for the association between SNPs and necrosis in the replicate dataset of 118 accessions.
