## Supplementary material for "Genetic basis of *Arabidopsis thaliana* responses to infection by naïve and adapted isolates of turnip mosaic virus": Figure 2 - figure supplement 2

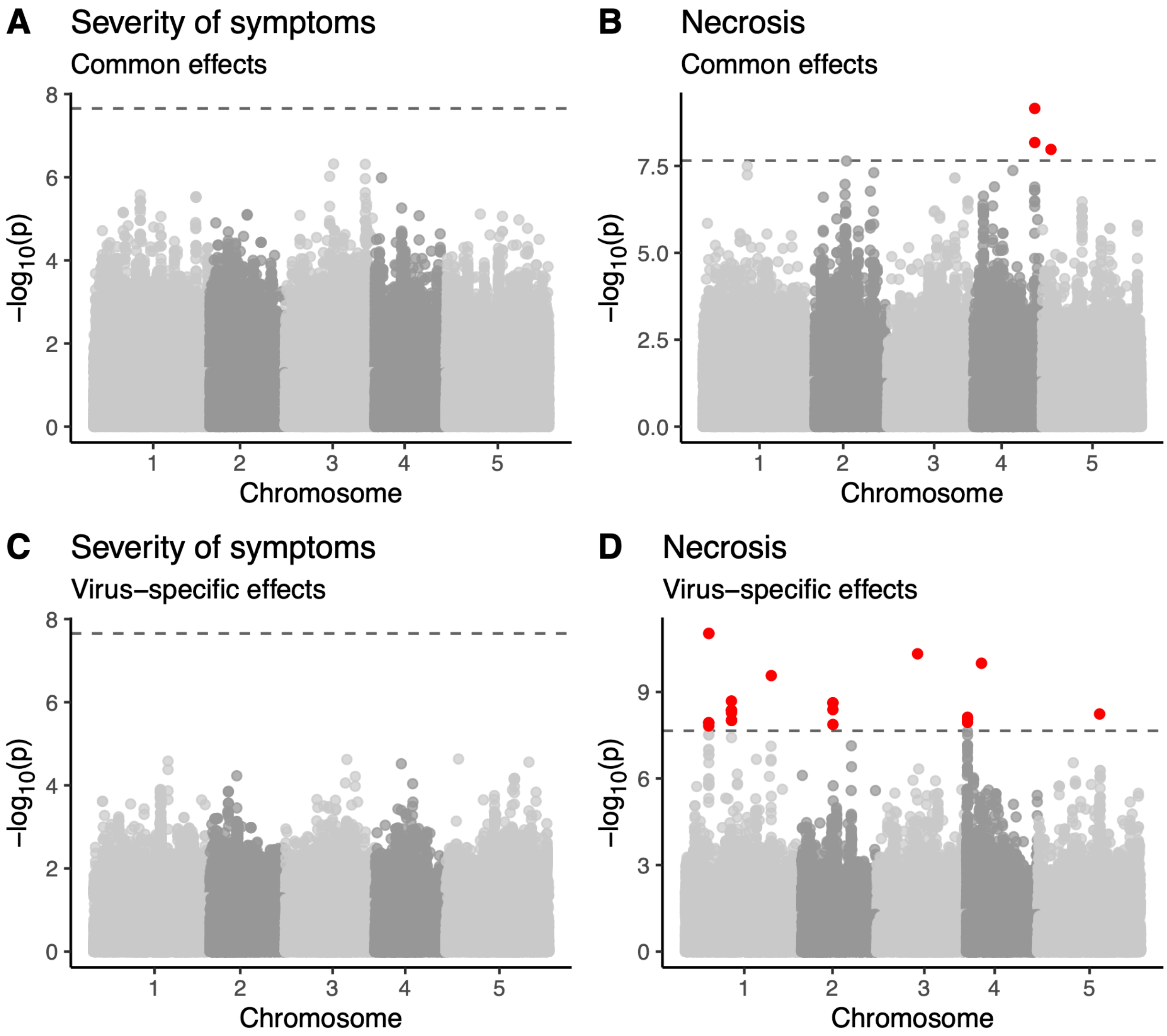


Figure S2: Manhattan plots for associations with severity and symptoms and necrosis in a model conditioned on the most strongly associated SNP (Chr2:5927469).
