## Supplementary figures and images for "Genetic basis of *Arabidopsis thaliana* responses to infection by naïve and adapted isolates of turnip mosaic virus"

### Figure 2 - figure supplement 3

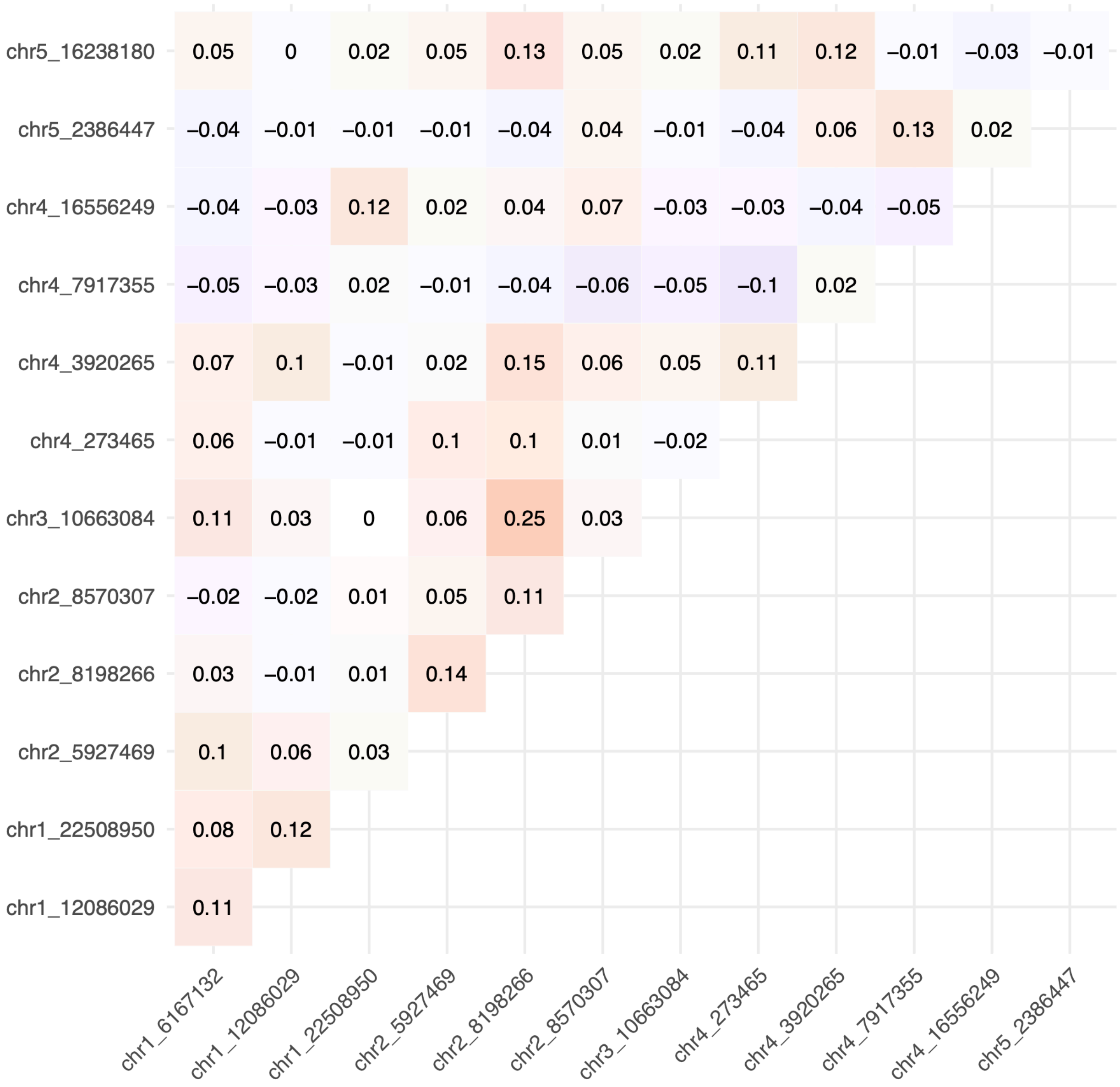


Figure S3: Linkage disequilibrium between SNPs associated with necrosis.
