## Supplementary material for "Genetic basis of *Arabidopsis thaliana* responses to infection by naïve and adapted isolates of turnip mosaic virus": Figure 2 - figure supplement 4

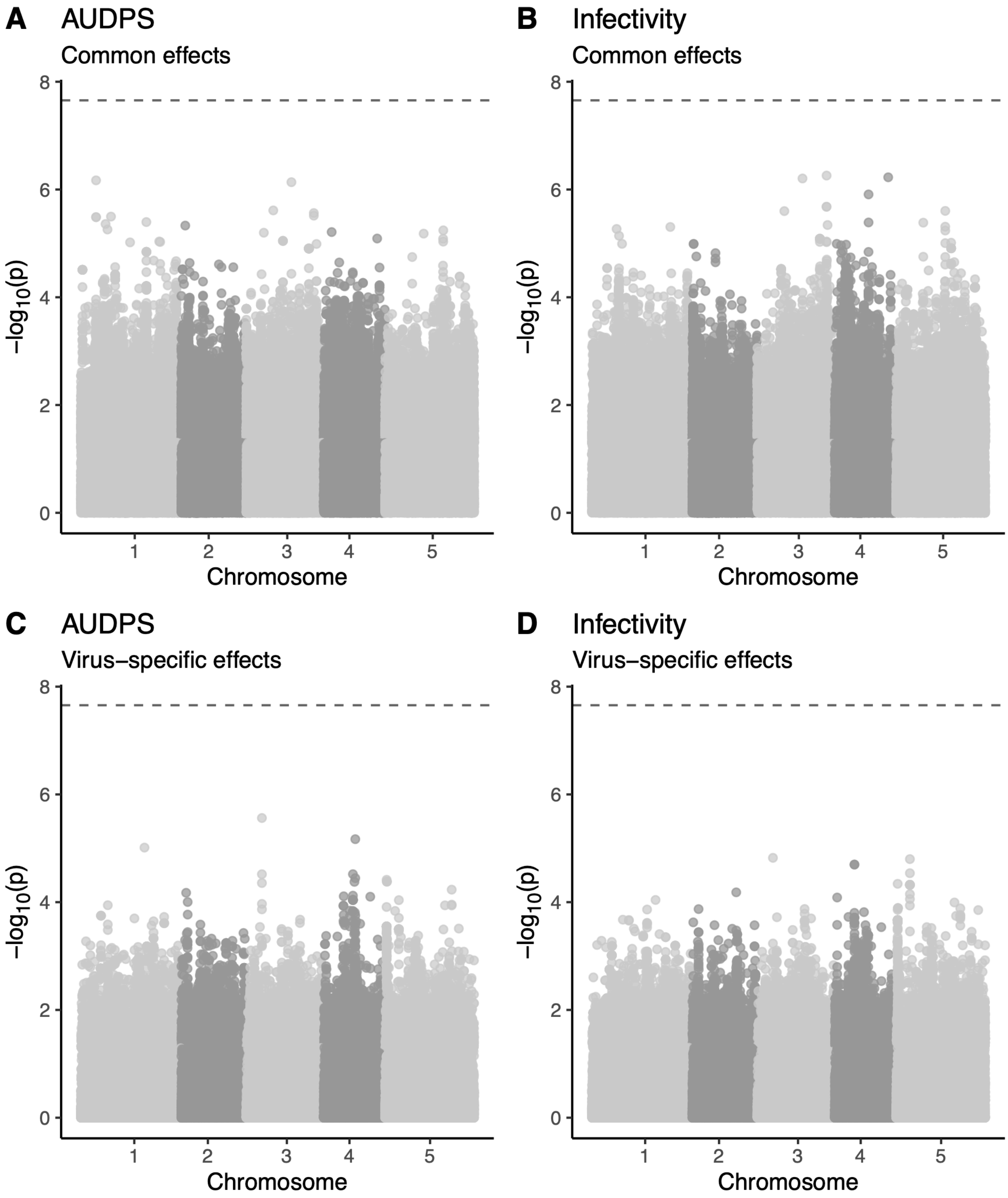


Figure S4: Manhattan plots for common and virus-specific associations with AUDPS and infectivity.
